# Systematics of a radiation of Neotropical suboscines (Aves: Thamnophilidae: *Epinecrophylla*)

**DOI:** 10.1101/2020.03.26.009746

**Authors:** Oscar Johnson, Jeffrey T. Howard, Robb T. Brumfield

## Abstract

The stipple-throated antwrens of the genus *Epinecrophylla* (Aves: Thamnophilidae) are represented by eight species primarily found in the lowlands of the Amazon Basin and the Guiana Shield. The genus has a long and convoluted taxonomic history, with many attempts made to address the taxonomy and systematics of the group. Here we employ massively parallel sequencing of thousands of ultraconserved elements (UCEs) to provide both the most comprehensive subspecies-level phylogeny of *Epinecrophylla* antwrens and the first population-level genetic analyses for most species in the genus. Most of our analyses are robust to a diversity of phylogenetic and population genetic methods, but we show that even with thousands of loci we are unable to confidently place the western Amazonian taxon *pyrrhonota*. We uncovered phylogenetic relationships between taxa and patterns of population structure that are discordant with both morphology and current taxonomy. For example, we found deep genetic breaks between taxa in the *ornata* group currently regarded as species, and in the *haematonota* and *leucophthalma* groups we found paraphyly at the species and subspecies levels, respectively. Our population genetics analyses showed extensive admixture between some taxa despite their deep genetic divergence. We present a revised taxonomy for the group, discuss the biogeographic patterns that we uncover, and suggest areas for further study.

## 1. Introduction

The stipple-throated antwrens of the genus *Epinecrophylla* (Isler et al. 2006; Aves: Thamnophilidae) are represented by 21 currently recognized taxa, eight of which are considered species (*E. fulviventris*, *E. ornata*, *E. erythrura*, *E. leucophthalma*, *E. gutturalis*, *E. amazonica*, *E. spodionota*, and *E. haematonota*; Figure 1). These species are primarily found in the lowlands of the Amazon Basin and the Guiana Shield, with one (*E. fulviventris*) found west of the Andes from Ecuador to Honduras (Clements et al., 2019; Zimmer and Isler, 2003). All species are small, insectivorous dead-leaf foraging specialists, typically found in pairs or small family groups in tropical upland forest (Remsen and Parker, 1984; Wiley, 1971). The genus reaches its greatest diversity in the western Amazon Basin, with up to three species broadly co-occurring in most regions, despite similar plumage and foraging behavior between species (Remsen and Parker, 1984; Zimmer and Isler, 2003).

**Figure 1.**
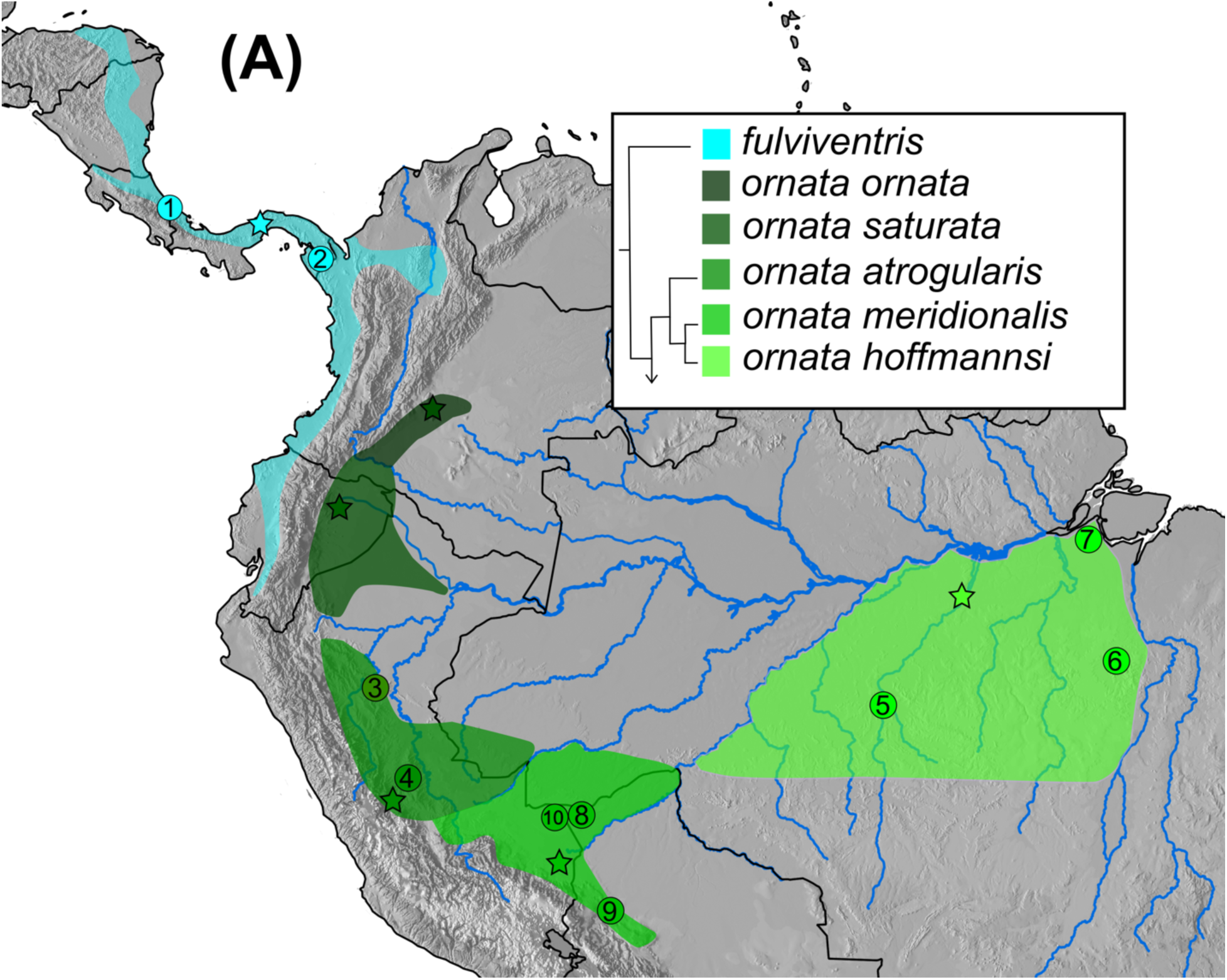
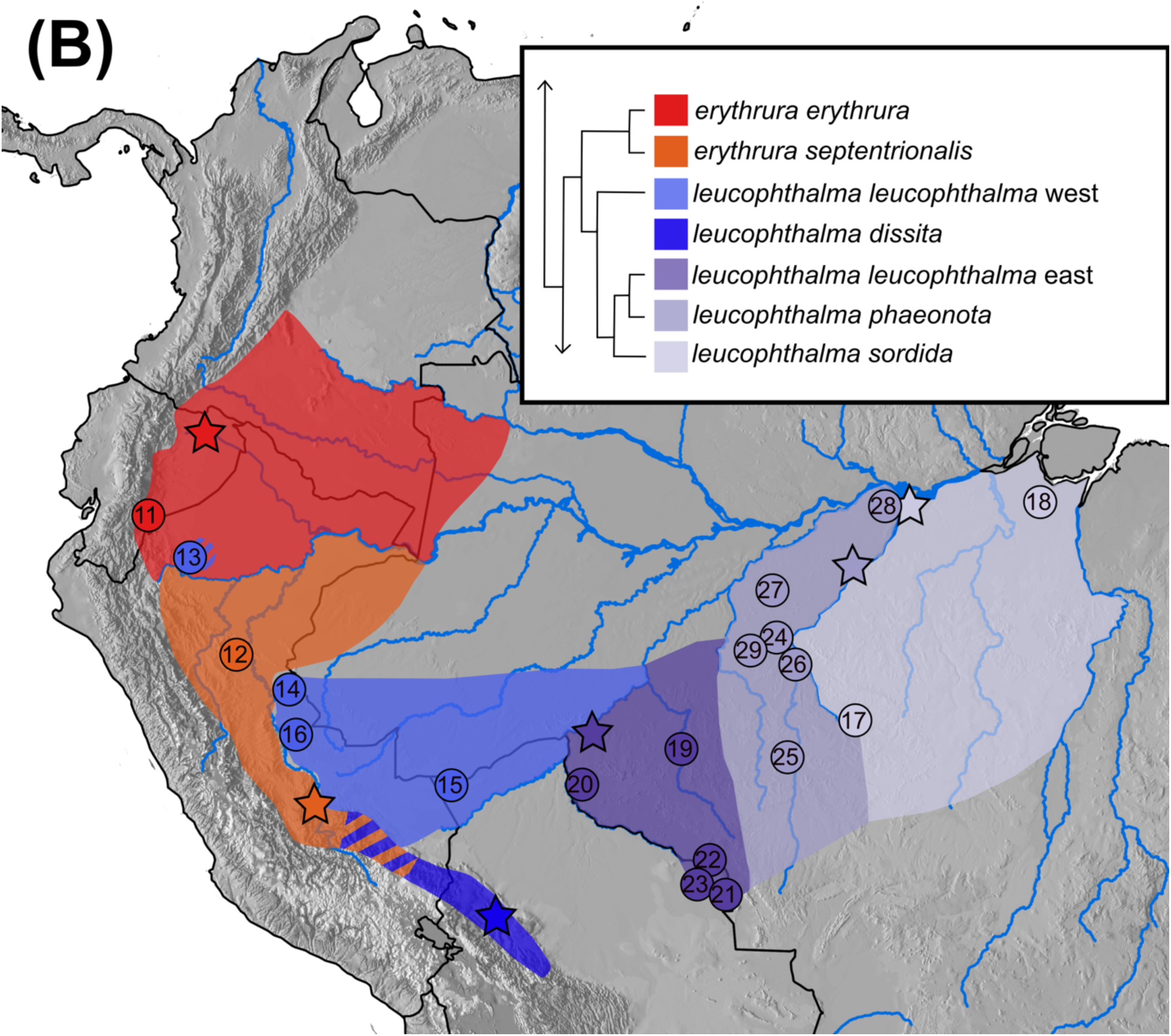
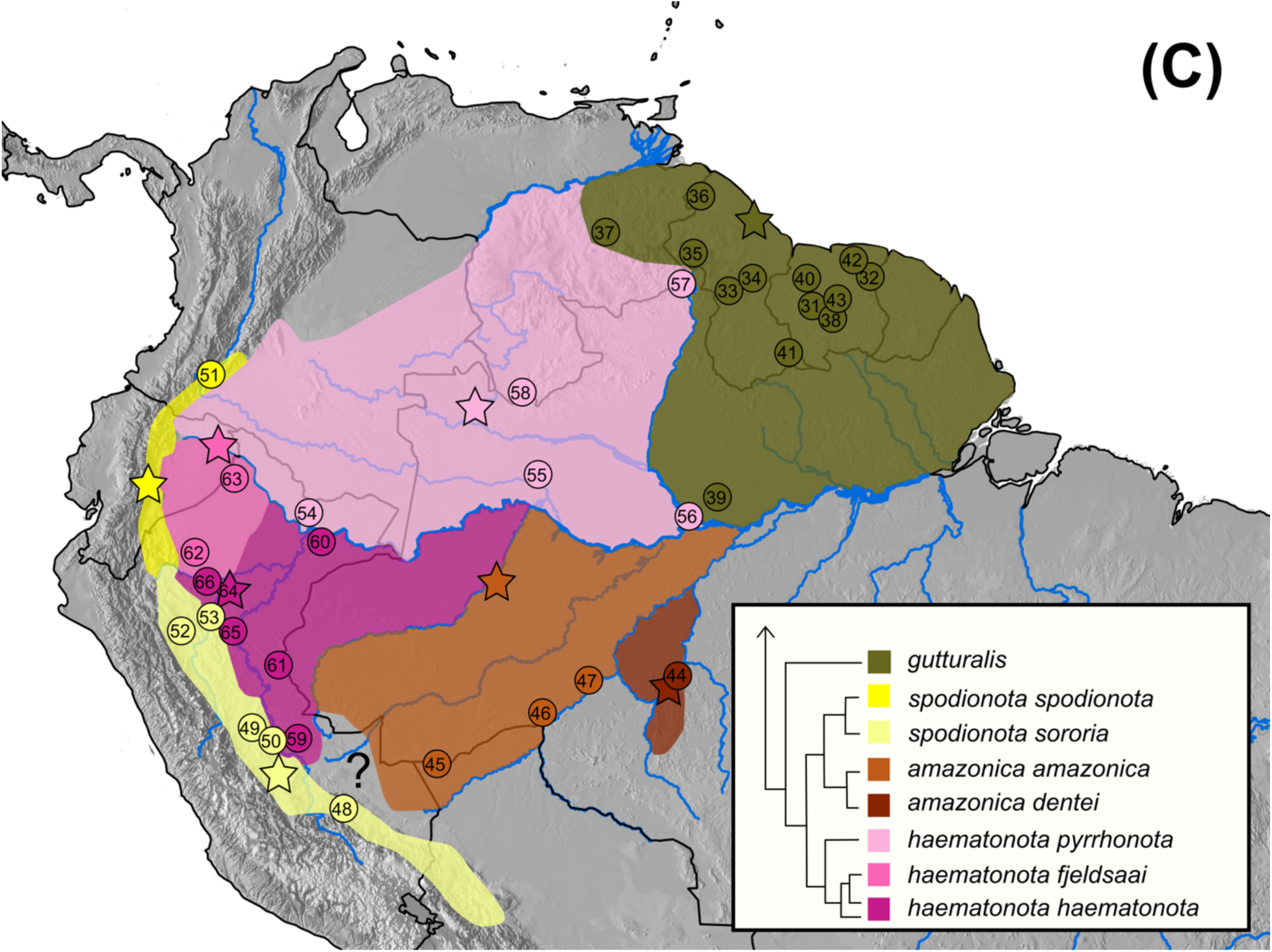
Maps showing taxon distributions, type localities, and sample localities used in this study. A) *Epinecrophylla fulviventris* and *ornata*. B) *E. erythrura* and *leucophthalma*. C) *E. gutturalis, pyrrhonota, dentei, amazonica, spodionota, haematonota,* and *fjeldsaai*. Country boundaries are shown in black. Major rivers are shown in blue. Locations sampled for this study are indicated with a number, corresponding to sample information in Table 1. Type localities, shown with a star, are based on Peters (1951) or type descriptions. Hashed regions indicate range overlap. Inset for each map shows a cladogram of relationships between each taxon based on the trees in Figure 2 and Figure S1H.

**Table 1.**
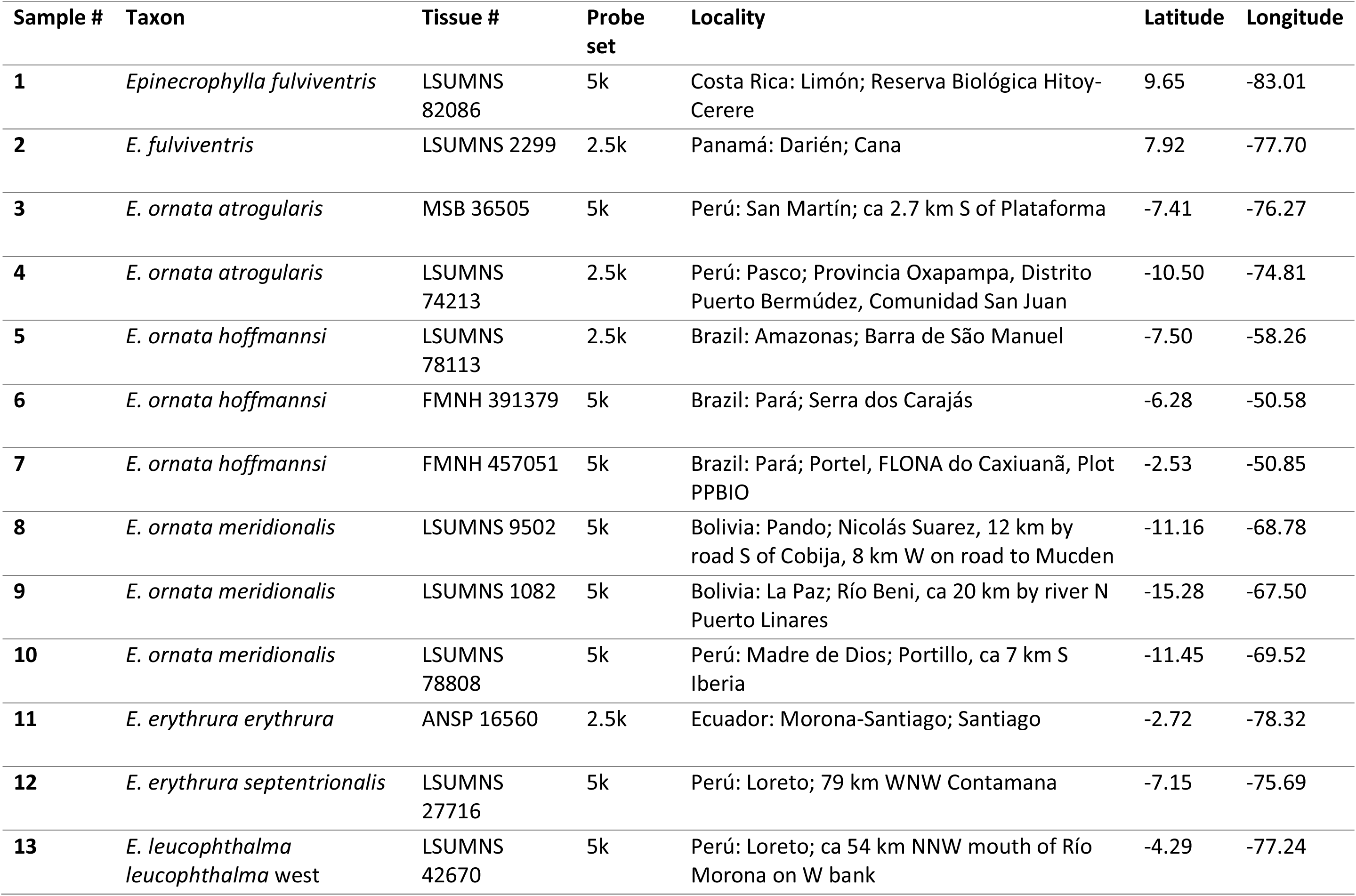
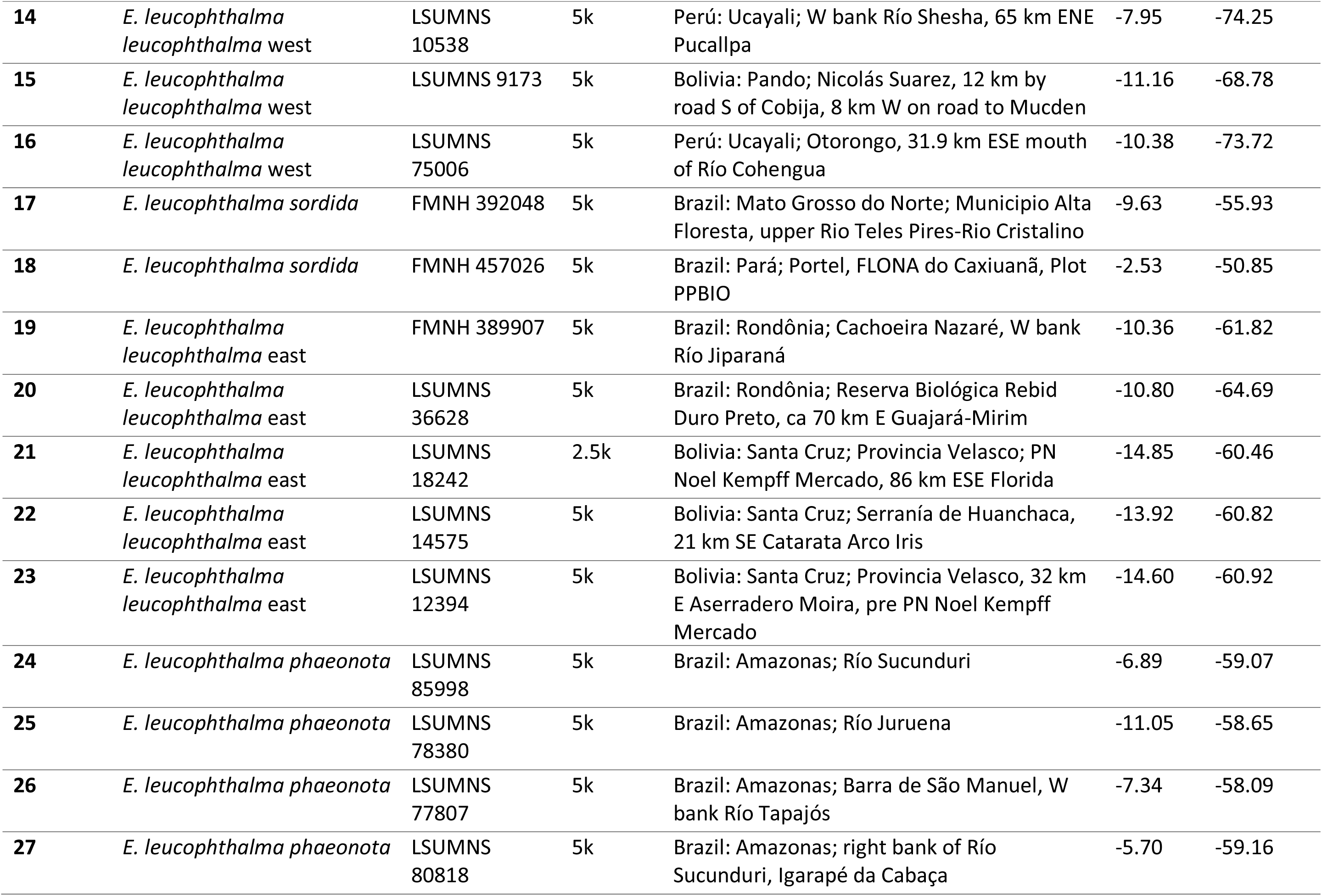
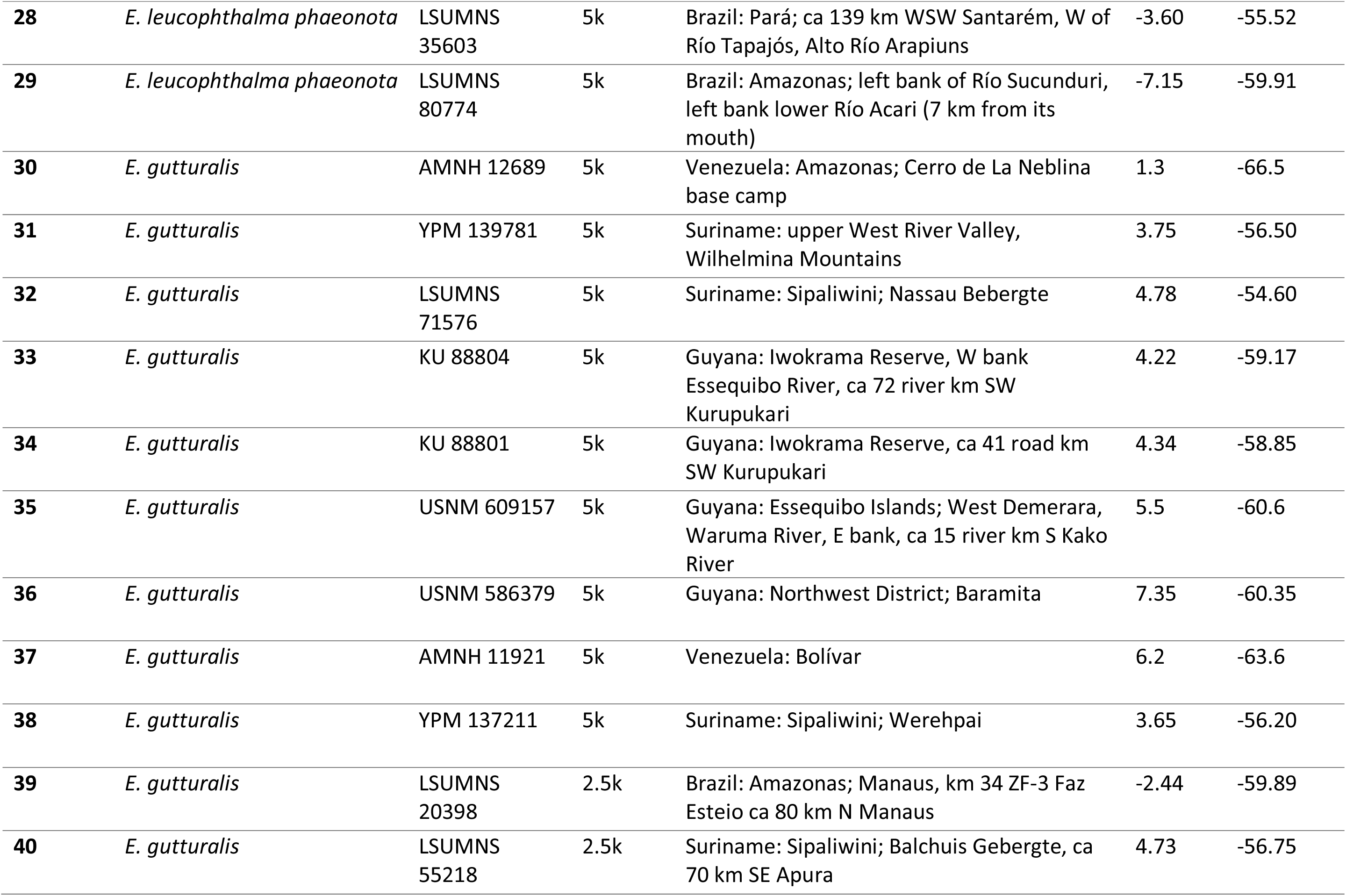
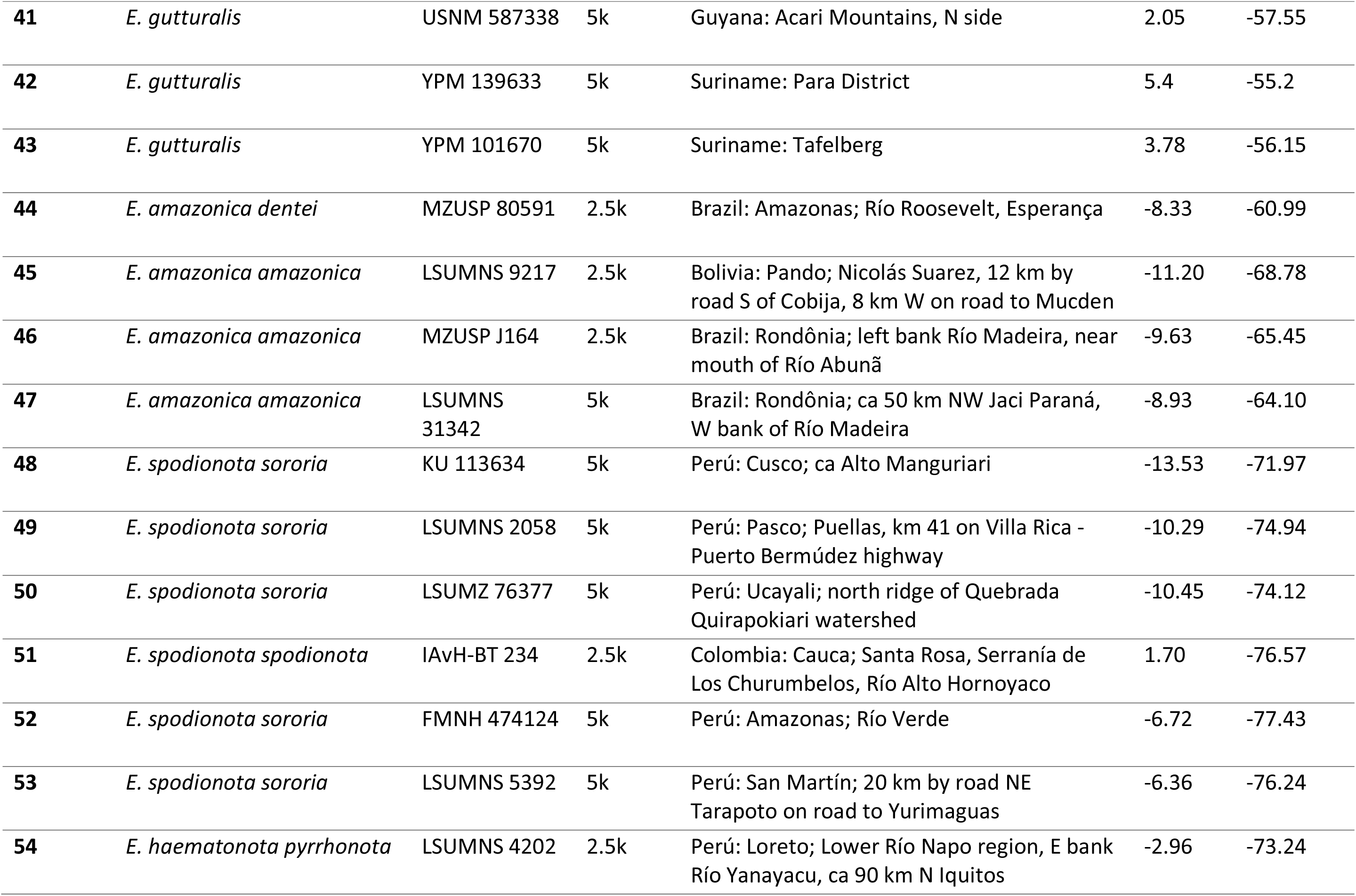
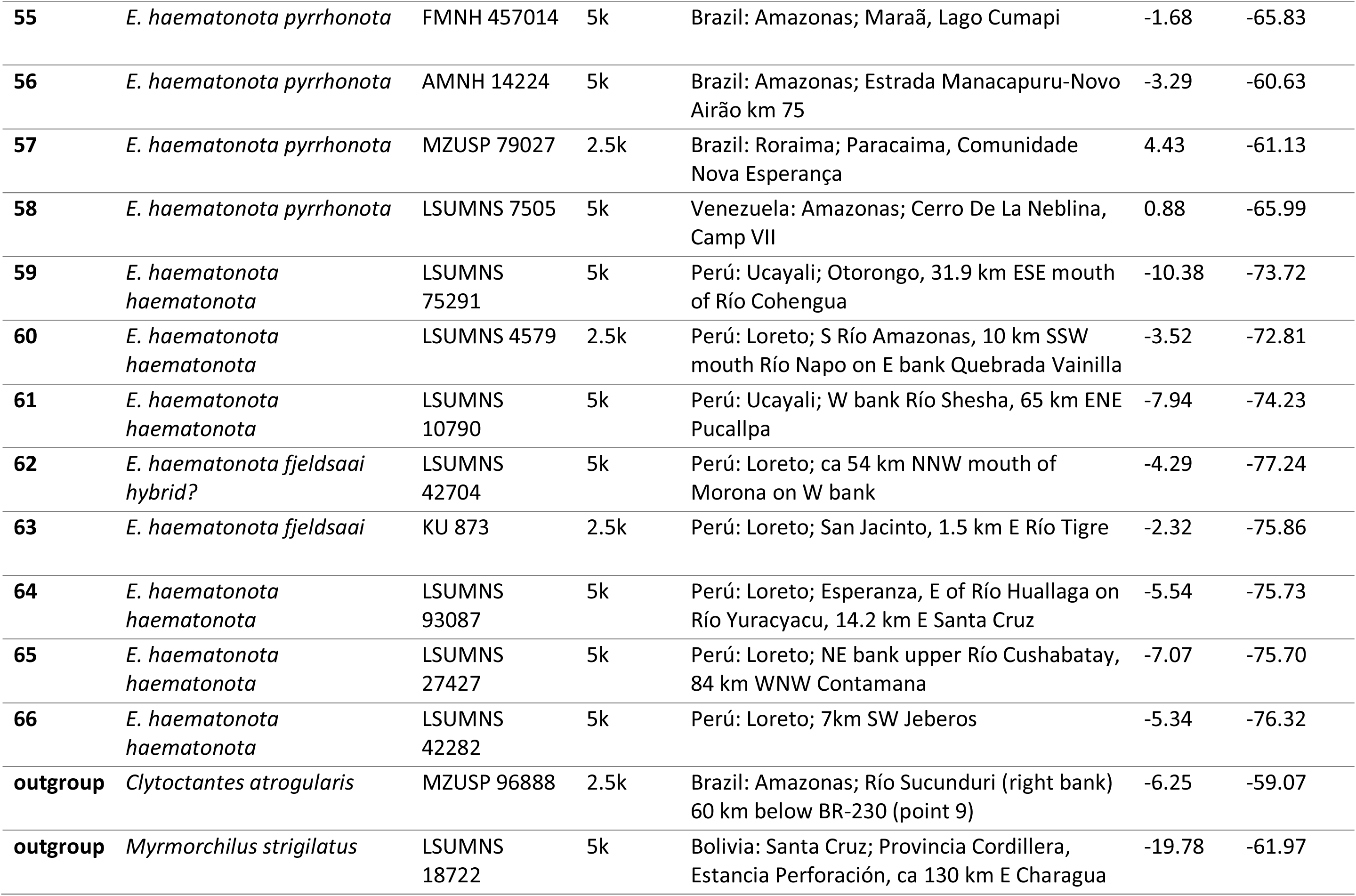
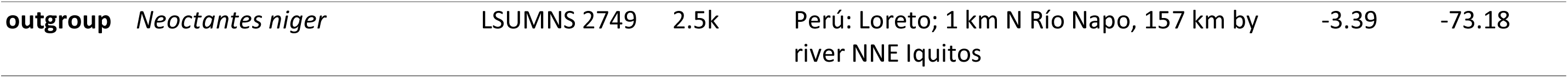
Localities for samples used in this project. Abbreviations are as follows: LSUMNS = Louisiana State University Museum of Natural Science, KU = University of Kansas Biodiversity Institute & Natural History Museum, AMNH = American Museum of Natural History, MZUSP = Museum of Zoology of the University of São Paulo, FMNH = Field Museum of Natural History, MSB = Museum of Southwestern Biology, USNM = Smithsonian National Museum of Natural History, YPM = Yale Peabody Museum. Probe set refers to the probe set used in sequencing. 5k = Tetrapods-UCE-5Kv1 probe set targeting 5,060 loci, and sequenced for this study. 2.5k = Tetrapods-UCE-2.5Kv1 probe set targeting 2,386 loci, sequences obtained from Harvey et al. (in press).

Multiple attempts have been made to resolve relationships in the genus with molecular data, with increasing numbers of loci and individuals used (Hackett and Rosenberg, 1990; Harvey et al., in press; Whitney et al., 2013). Long considered to be in the genus *Myrmotherula*, early molecular work using protein electrophoresis provided the first indication that the stipple-throated antwren complex was not a close relative of other *Myrmotherula* antwrens (Hackett and Rosenberg, 1990). This was further supported with Sanger sequencing of mitochondrial and nuclear loci (Bravo et al., 2014; Brumfield et al., 2007; Irestedt et al. 2004), with the studies finding that *Epinecrophylla* was most closely related to bushbirds in the genera *Neoctantes* and *Clytoctantes*. This work led to the naming of a new genus for the group, *Epinecrophylla* (Isler et al., 2006), with *E. haematonota* as the type species. Some authorities changed the common names of *Epinecrophylla* antwrens to stipplethroats (Clements et al., 2019; Remsen et al., 2019) to reflect this taxonomic rearrangement.

### 1.1 History of taxa within Epinecrophylla

The species-level taxonomy of the genus has undergone considerable rearrangement through history (Cory and Hellmayr, 1924; Isler and Whitney, 2018; Peters, 1951; Whitney et al., 2013; Zimmer, 1932a; 1932b), particularly in the *haematonota* and *leucophthalma* groups. Early authors (e.g. Cory and Hellmayr, 1924) considered *E. haematonota* to include as subspecies the taxa *pyrrhonota* and *amazonica* and placed both *spodionota* and *sororia* as subspecies of *E. leucophthalma*, largely based on back color (rufous in the former three taxa, brown in the latter three). Using this same reasoning, Todd (1927), when describing the rufous-backed form *phaeonota*, treated it as a subspecies of *E. haematonota*, but considered *E. amazonica* a species distinct from all other forms. Zimmer (1932a), however, noted that back color may not be a valid species-level character and transferred *amazonica, spodionota*, and *sororia* to *E. haematonota*, and *phaeonota* to *E. leucophthalma*. Zimmer (1932a) suggested the possibility of species rank for the rufous-backed taxon *phaeonota*, but also noted intermediate individuals between it and the adjacent brown-backed taxa *leucophthalma* and *sordida*. This treatment was maintained by most authors (e.g. Meyer de Schauensee, 1970; Peters, 1951) until Parker and Remsen (1987) recognized *E. spodionota* (including *sororia*) as a separate species. This taxonomic treatment was augmented by the recent discovery of two range-restricted taxa in the group; *E. fjeldsaai* of eastern Ecuador (Krabbe et al., 1999) and *E. dentei* of the Aripuanã-Machado region of Brazil (Whitney et al., 2013), each described as a new species. In describing *E. dentei* Whitney et al. (2013) also estimated a mitochondrial phylogeny of the genus, including samples of most taxa, in which they found *fjeldsaai* was phylogenetically embedded within *haematonota*. Based on that work and the mitochondrial distance between taxa, Remsen et al. (2019) separated *E. haematonota* into four species: *E. fjeldsaai* (based on morphology)*, E. pyrrhonota*, *E. haematonota*, and *E. amazonica* (including *dentei*), whereas other authors united *pyrrhonota*, *amazonica*, and *dentei* under *E. haematonota* while maintaining *E. fjeldsaai* as a distinct species (Dickinson and Christidis, 2014). Since then, Isler and Whitney (2018) conducted a thorough analysis of the vocalizations of *haematonota*, *fjeldsaai*, and *pyrrhonota* in which they found no vocal differences among the three taxa, leading to the recognition of the latter two taxa as subspecies of the former (Remsen et al., 2019).

Within *E. ornata*, the gray-backed Peruvian taxon *atrogularis* was long considered a separate species, leaving the rufous-backed forms *saturata* and *hoffmannsi* as subspecies of *E. ornata* (Cory and Hellmayr, 1924). This treatment was maintained until Zimmer (1932b) described the gray-backed *meridionalis* as a subspecies and united all five taxa in the group under the species *E. ornata*. This is the current treatment of most recent authors (Clements et al., 2019; Dickinson and Christidis, 2014; Remsen et al., 2019), although del Hoyo et al. (2019) consider *E. hoffmannsi* a species separate from the rest of *E. ornata* based primarily on the female plumage and minor vocal differences.

The taxonomy of the remainder of the genus has remained rather more stable through time, with the three other species *E. fulviventris*, *E. gutturalis,* and *E. erythrura*, all largely considered independent species by most authors. *E. erythrura* and *E. leucophthalma* are currently considered polytypic, while the four taxa described in *E. fulviventris* are generally considered synonyms of the nominate subspecies (Cory and Hellmayr, 1924; Zimmer and Isler, 2003). We here follow the taxonomy of the South American Classification Committee (Remsen et al., 2019) and make taxonomic recommendations in light of the Biological Species Concept (de Queiroz, 2007; Mayr, 1942).

Much of the previous molecular work in *Epinecrophylla* has relied upon mitochondrial sequence data, although a recent phylogenomic study of all suboscine passerine birds included 1-2 samples of each species of *Epinecrophylla* using sequence capture of ultraconserved elements (UCEs) and recovered a well-resolved topology for the genus (Harvey et al., in press). Here we expand on the previous genetic work in the genus, addressing the systematics of the group with both next-generation sequencing of thousands of nuclear loci and draft mitochondrial genomes, and population-level sampling of most taxa. *Epinecrophylla* provide a unique system in which to study speciation in the Amazon Basin due to their high species diversity, documented phenotypic hybrid zones, and multiple broadly sympatric species. Our expanded sampling both of individuals and loci provides the most in-depth view of the evolutionary history, species limits, population structure, and introgression between taxa in this group.

## 2. Methods

### 2.1. Sampling

We obtained a total of 66 *Epinecrophylla* and three outgroup samples representing 18 of the 21 widely recognized taxa in the genus and all currently recognized species. Missing ingroup taxa are *E. o. ornata*, *E. o. saturata*, and *E. leucophthalma dissita*. The outgroup species we used are *Myrmorchilus strigilatus*, *Neoctantes niger*, and *Clytoctantes atrogularis*. When available, we obtained samples from across the geographic range of each *Epinecrophylla* taxon, with one sample chosen per geographic locality. Fifty-three tissue samples were obtained from vouchered specimens housed at museums in the United States, with sequence data for the remaining 16 samples obtained from Harvey et al. (in press; Table 1).

We extracted total DNA from the 53 tissue samples using ca. 25 mg of pectoral muscle with a Qiagen DNeasy Blood and Tissue Kit (Qiagen; Hilden, Germany) and quantified DNA concentration using a Qubit 2.0 fluorometer (Life Technologies Corporation; Carlsbad, CA). Samples were standardized to 10 ng/uL. We then sheared samples to approximately 600 base pair (bp) fragments with an Episonic 1100 bioprocessor (EpiGentek; Farmingdale, NY) and assessed fragment length using a High Sensitivity DNA Assay on an Agilent 2100 Bioanalyzer (Agilent Technologies; Santa Clara, CA). We generated DNA libraries using a KAPA Biosystems Hyper Prep kit (Wilmington, Massachusetts, USA) and enriched UCEs using a set of 5,742 probes that target 5,060 loci in vertebrates (“Tetrapods-UCE-5Kv1”; uce-5k-probes.fasta) following the protocol of Faircloth et al. (2012). Enriched samples were pooled at equimolar ratios and paired-end sequencing was conducted on one lane of a HiSeq 3000 sequencer at Oklahoma Medical Research Foundation Clinical Genomics Center (OMRF; Oklahoma City, Oklahoma, USA). The sequencing lane contained DNA libraries used in other projects. The 16 samples obtained from Harvey et al. (in press) were enriched using a custom probe set consisting of 2,500 vertebrate UCEs and 96 exons.

### 2.2. Contig assembly

OMRF demultiplexed sequence reads using custom scripts. We trimmed raw reads of adapter contamination and low-quality bases using illumiprocessor (Faircloth, 2013) and trimmomatic (Bolger et al., 2014) with default settings. We then subsampled all read files to 2 million reads per individual to decrease computation time for contig assembly and to normalize assemblies across samples. Read data were assembled with Itero (https://github.com/faircloth-lab/itero). Because samples were sequenced with two different probe sets, we opted to match contigs to the “Tetrapods-UCE-2.5Kv1” (uce-2.5k-probes.fasta) probe set which consists of 2,560 baits targeting 2,386 UCE loci, and is a subset of the other probe sets used in sequencing. For the samples sequenced with the “Tetrapods-UCE-5Kv1” probe set, we also separately matched assembled contigs to this probe set.

### 2.3. Sample identification and locus filtering

To confirm the identifications of samples we used the Phyluce 1.6.7 (Faircloth, 2015) tool *match_contigs_to_barcodes* to match contigs from each sample to a mitochondrial Cytochrome c oxidase subunit I (COI) barcode sequence of *Epinecrophylla pyrrhonota* obtained from GenBank (JN801852.1) and map those contigs against the Barcode of Life Database (BOLD; Ratnasingham and Hebert, 2007). We then used the Phyluce 1.6.7 (Faircloth, 2015) tool *get_trinity_coverage* to calculate per-locus read coverage for all contigs matching either UCE and mitochondrial loci. Three samples contained mitochondrial loci with high coverage (>30x) that matched the incorrect species in BOLD, suggesting either sample misidentification or high levels of contamination, and were eliminated from our dataset (Table S1). Nine additional samples contained high-coverage mitochondrial contigs matching the expected species in BOLD, but with a small number of low-coverage mitochondrial contigs matching the incorrect species. We used the maximum coverage of 8.05x of these potentially contaminated low-coverage mitochondrial contigs as a filter and removed all UCE contigs across all samples that had an average read depth below this threshold.

### 2.4. Mitochondrial genome assembly

We used off-target reads from the UCE sequencing to assemble draft mitochondrial genomes. We assembled mitochondrial genomes in MITObim 1.9 (Hanh et al., 2013), which is a Perl wrapper for MIRA 4.0.2 (Chevreux et al., 1999), using as a reference the complete mitochondrial genome of *Myrmoderus loricatus* (G. Bravo, unpublished data) and the --*quick* option. We annotated the assembled mitochondrial genomes using the MITOchondrial genome annotation Server (MITOS) 2 (Bernt et al., 2013) and aligned the 13 mitochondrial protein coding genes in MAFFT (Katoh et al., 2002) implemented in Geneious 10.2.3 (https://www.geneious.com) to create a final partitioned draft mitogenome alignment.

### 2.5. Nuclear locus phasing, alignment, and SNP calling

To phase UCE loci we selected as a reference the individual from our sampling that contained the greatest number of UCE loci after filtering; *Epinecrophylla leucophthalma* LSUMNS 42670. We phased UCE loci using the *seqcap_pop* pipeline (https://github.com/mgharvey/seqcap_pop; Faircloth, 2015; Harvey et al., 2016) to obtain a Single Nucleotide Polymorphism (SNP) dataset and followed Andermann et al. (2019) to obtained phased alignments. The *seqcap_pop* pipeline utilizes tools from the Phyluce package (Faircloth, 2015), SAMtools 0.1.19 (Li et al., 2009), Picard (Broad Institute, Cambridge, MA), BWA 0.7.17 (Li and Durbin, 2009), and GATK 3.3.0 (McKenna et al., 2010) to process next-generation sequence data for population-level genetic analyses. Briefly, *seqcap_pop* maps sequencing reads to the reference individual to obtain a pileup, adds read groups and marks PCR duplicate reads for each individual, merges bam files within each species, calls indels and single-nucleotide polymorphisms on merged bam files, and phases high-quality indels and SNPs to produce vcf files of phased SNPs. We further filtered this dataset using vcftools 0.1.16 (Danecek et al., 2011) to remove SNPs with quality scores less than 30 and read depth less than 5.5, those with greater than 75% missing data, restricted to bi-allelic loci, and removed indels. We refer to this dataset as the “linked SNP dataset”, as it contains multiple SNPs per locus. We then sampled at random one SNP per UCE locus to obtain a final dataset of putatively unlinked SNPs, which we refer to as the “unlinked SNP dataset”. To obtain phased alignments we used Phyluce 1.6.7 (Faircloth, 2015) to phase UCE loci following Andermann et al. (2019), phasing data by mapping reads against the reference individual using the Phyluce tools *snp_bwa_align* and *snp_phase_uces*. This pipeline maps raw reads against contigs of a reference individual using BWA 0.7.17 (Li and Durbin, 2009), and then sorts and phases alleles in SAMtools 0.1.19 (Li and Durbin, 2009) and Picard (Broad Institute, Cambridge, MA). We used MAFFT 7.130b (Katoh and Standley, 2013) in the Phyluce tool *align_seqcap_align* to align and edge-trim the contigs output by this pipeline, treating the two alleles as separate individuals and allowing ambiguous sites in alignments. We produced a final alignment using the Phyluce 1.6.7 (Faircloth, 2015) tool *get_only_loci_with_min_taxa* to produce a 75% complete data matrix. This tool calculates the data matrix completeness as the percentage of individuals in the dataset with sequence data for each locus.

To investigate fine-scale patterns of population structure within each species we called SNPs within each species or species complex to obtain an additional six SNP datasets. We grouped samples based on the clades in the Exabayes phylogeny estimated from the 75% complete UCE data matrix (see section 2.6). Three clades corresponded to species (*ornata*, *leucophthalma*, and *gutturalis*) and a fourth to a set of closely related taxa that have undergone considerable taxonomic rearrangement through history (*dentei*, *amazonica*, *spodionota*, *sororia*, *pyrrhonota*, *haematonota*, and *fjeldsaai*). This latter clade is hereafter referred to as the “*haematonota s.l*.” clade. Within the *haematonota s.l*. clade we additionally subdivided taxa into two clades for SNP calling: one containing *dentei*, *amazonica*, *spodionota*, and *sororia* (hereafter the “*amazonica* + *spodionota*” clade), and the other containing *pyrrhonota*, *haematonota*, and *fjeldsaai* (hereafter the “*haematonota* + *pyrrhonota*” clade). For each dataset we selected as a reference the individual with the highest number of assembled contigs after filtering (Supplemental Table 4) and repeated the *seqcap_pop* pipeline described above.

### 2.6. Phylogenetic estimation

From the 2,386 locus UCE dataset we estimated a phylogenetic tree with all samples using a Bayesian analysis in Exabayes 1.5 (Aberer et al., 2014) using the 75% complete concatenated phased alignment. We conducted 4 independent runs for 2 million generations each, discarding the first 25% of trees as burn-in. After checking in Tracer 1.7.1 (Rambaut et al., 2018) that samples had converged based on ESS values greater than 200, we generated an extended majority-rule consensus tree using the topologies from the four independent runs.

We used this topology to estimate a time-calibrated phylogenetic tree in BEAST 2.5.2 (Bouckaert et al., 2019). For this analysis we took the tree estimated in Exabayes and trimmed it to include only the samples for which we obtained draft mitochondrial genomes (the Exabayes UCE phylogeny containing all samples is available in Figure S1), and one allele per individual.

We then made this tree ultrametric using the *chronos* function in ape 5.2 (Paradis and Schliep, 2018) and used the resulting topology as a constraint tree through the entire BEAST analysis. No fossils are available for *Epinecrophylla* or its close relatives to allow for a fossil calibration of our phylogenetic tree. We therefore used three combinations of mitochondrial alignments and substitution rates to estimate the branch lengths for this topology. We first used the concatenated alignment of the 13 mitochondrial protein-coding genes and the widely used mitochondrial cytochrome B (CytB) 2.1% substitution rate obtained from Hawaiian honeycreepers (Lerner et al., 2011; substitution rate: 0.01105 substitutions/site/Myr, 95% Confidence Interval [CI]: 0.00425–0.01785). The second method used the alignment of cytochrome B (CytB) extracted from the draft mitochondrial genomes and the same CytB substitution rate. For the final analysis we analyzed the third codon position of CytB and used a mass-calibrated substitution rate following (Nabholz et al., 2016) using the third codon position and their calibration set 2 (substitution rate: 0.0208 substitutions/site/Myr, 95% CI: 0.0170– 0.0263). Body mass estimates were obtained from specimens at the Louisiana State University Museum of Natural Science (mean = 9.30 g, SD = 1.18, n = 229), representing all species in the genus. We used the date estimate from a dated phylogeny of all suboscine passerines to constrain the root of our phylogeny (Harvey et al., in press), and set *E. fulviventris* as the outgroup, with the date of 9.28 Ma (95% CI: 8.60–11.07 Ma) as the prior on the root node. For each analysis we used the GTR + γ model of rate variation and a proportion of invariant sites, a log-normal relaxed clock, a yule model on the tree, and default settings for other priors. We ran each analysis for 100 million generations, sampling every 10,000, and a burn-in of 10%. We checked in Tracer 1.7.1 (Rambaut et al., 2018) that all parameters reached convergence with ESS values over 200. Both analyses using the CytB alignment were unable to converge and had multiple priors with ESS values well below 200, indicating inappropriate priors and/or insufficient generations. The analysis using the draft mitochondrial genome alignment, however, converged with ESS values over 200 for all parameters. For the mitogenome analysis we calculated a maximum clade credibility (MCC) tree in TreeAnnotator 2.5.1 from the posterior of trees, implemented in BEAST 2.5.2 (Bouckaert et al., 2019), and visualized the resulting tree in FigTree 1.4.4 (Rambaut, 2009).

We used SNAPP 1.4.2 (Bryant et al., 2012) implemented in BEAST 2.5.2 (Bouckaert et al., 2019) to calculate a species tree directly from SNP data in a full-coalescent analysis. This site-based method has the advantage of bypassing gene tree estimation and minimizing error due to the low information content of individual UCE loci. Initial runs using all samples in our dataset were unable to converge in a reasonable amount of time, either with individuals as separate tips or with individuals pooled to tips by clade identified from the Exabayes 75% UCE phylogeny. Therefore, we called SNPs again using two sample-filtering and locus-filtering strategies, all with *Myrmorchilus strigilatus* as an outgroup; the first using up to two samples from each clade identified in the Exabayes 75% phylogeny for a total of 19 samples, and the second using one sample from each widely recognized species in the genus for a total of 11 samples. In both datasets we used the individual from each taxon that had the greatest number of loci recovered to maximize the number of SNPs recovered and allowed 5% missing data.

Otherwise we followed the *seqcap_pop* and SNP filtering pipeline described in section 2.5. After SNP filtering we selected at random one SNP per locus to minimize issues with linkage within a locus. In all runs we calculated the mutation rates from the data and used default priors. We ran all analyses for 2 million generations, storing every 1,000 generations, and a burn-in of 10%, and checked that the run converged in Tracer 1.7.1 (Rambaut et al., 2018) based on ESS values over 200. From the posterior of species trees we calculated a maximum clade credibility (MCC) tree in TreeAnnotator 2.5.1 implemented in BEAST 2.5.2 (Bouckaert et al., 2019). We used Densitree 2.0.1 (Bouckaert, 2010) to visualize the posterior tree set of the SNAPP runs and FigTree 1.4.4 (Rambaut, 2009) to visualize the MCC tree.

In addition to the analyses outlined above, we conducted a variety of phylogenetic analyses, each with its own assumptions, strengths, and weaknesses relative to treating sources of phylogenetic variation. Details and results for these analyses are available in the Supplementary Materials.

### 2.7. Population genetics and introgression

In addition to our phylogenetic analyses we utilized SNPs to investigate patterns of population-level genetic structure and also introgression within and between clades. We used STRUCTURE (Pritchard et al., 2000) and Discriminant Analysis of Principal Components (DAPC) to analyze patterns of population structure within each clade, and implemented each analysis on all six clade-level SNP datasets described above. For STRUCTURE we analyzed the linked SNP datasets and implemented the *linked* model, providing the distance in base pairs between SNPs within each locus, and ran analyses for 2 million generations, discarding the first 50,000 as burn-in. We ran 10 replicate analyses for each value of K from one to 10, or until the likelihood value of K decreased significantly. We selected the best K value based on the ΔK method of Evanno (Evanno et al., 2005), implemented in STRUCTURE HARVESTER (Earl and vonHoldt, 2012).

DAPC uses sequential k-means clustering of principal components to infer the number of genetic clusters in a dataset. We conducted a DAPC analysis in *adegenet* (Jombart and Ahmed, 2011), following the recommendations of Jombart et al. (2010), and selected the best number of clusters based on the lowest Bayesian Information Criterion (BIC) score. In addition, we conducted a Principal Components Analysis (PCA) analysis, with samples coded by DAPC group assignments. Although BIC scores for the *haematonota s.l.* clade indicated that K values greater than 2 had a worse fit to the data than K=2, we calculated DAPC group assignments for K values from 3–5 to investigate finer-scale patterns of population structure, due to the greater number of described taxa in this clade.

We calculated pairwise distance estimates between all taxa in the group from both the mitochondrial and nuclear DNA data. For the mitochondrial distances we used the alignment of the 13 mitochondrial protein-coding genes. On a neighbor-joining tree reconstructed from the raw *p*-distance matrix in PAUP* 4.0 (Swofford, 1999), we estimated the proportion of invariant sites (0.590355) and the gamma shape parameter (1.82626). These values were then fixed for calculations of a distance matrix under the GTR + γ + I finite-sites substitution model. For the nuclear data we estimated the weighted fixation index (F_st_) between each pair of taxa using the method of Weir and Cockerham (1984) implemented in vcftools 0.1.16 (Danecek et al., 2011) using the unlinked SNP dataset. For all calculations we also report the average within-taxon distance estimates as a measure of intra-specific genetic structure.

## 3. Results

### 3.1. Sequencing results and sample identification

Illumina sequencing of UCEs resulted in an average of 3.8 million reads per individual, and an average read length of 130 bp after trimming. After removing potentially contaminated or misidentified samples our dataset contained 63 *Epinecrophylla* samples and two outgroups. Including the three potentially contaminated *Epinecrophylla* samples (based on BOLD results) in a phylogenetic tree estimated in RAxML 8.2.12 (Stamatakis, 2014), two grouped with the correct taxon but sat on abnormally long terminal branches, suggesting contamination, and a third grouped with the outgroup samples, suggesting sample misidentification (Figure S1H).

After assembly and locus filtering we obtained an average of 2,195 UCE loci per sample (range 1,234—2,306 loci), with a mean locus length of 652 bp (range 234–1,283 bp) and mean read depth of UCE loci of 22.5x (SD: 43.0x). Missing data had a strong effect on the number of UCE loci retained in the alignment, and the alignment that including no missing data contained 330 UCE loci and was not analyzed further. The 95% complete phased alignment contained 1,659 UCE loci and an aligned matrix of 1,140,275 bp and the 75% complete phased alignment contained 2,149 UCE loci and an aligned matrix of 1,401,699 bp.

We obtained partial or complete mitochondrial genomes for 59 ingroup samples and both outgroups, including at least one sample per species (Table S2). Three samples, including one of the outgroups, contained greater than 40% missing data and were removed from the analysis (Table S2). The average mitochondrial genome size was 17,253 bp (range 16,017–17, 930 bp) and had a mean read depth of 304.4x (SD: 780.0x). The aligned dataset of 56 individuals using the 13 mitochondrial protein-coding genes was 11,488 bp in length (range 9,921–11,396 bp), contained a total of 635,429 bp, and 4.8% missing data.

### 3.2. Phylogenetic estimation

From the nuclear UCE data, we recovered a phylogeny with strong support for relationships among taxa (Figure 2, Figure 3, Figure S1). The deepest split in the tree occurred across the Andes, dividing *E. fulviventris* from the remainder of the genus. Although our sampling included just two samples of *E. fulviventris*, one of which (sample 1) is from a population occasionally separated as the subspecies *costaricensis* (del Hoyo et al., 2019; Todd, 1927), our phylogenies indicated a relatively shallow divergence between the two samples (Figure S1). The next split separated *E. ornata* from the remaining taxa (Figure 1A, Figure 2).

**Figure 2.**
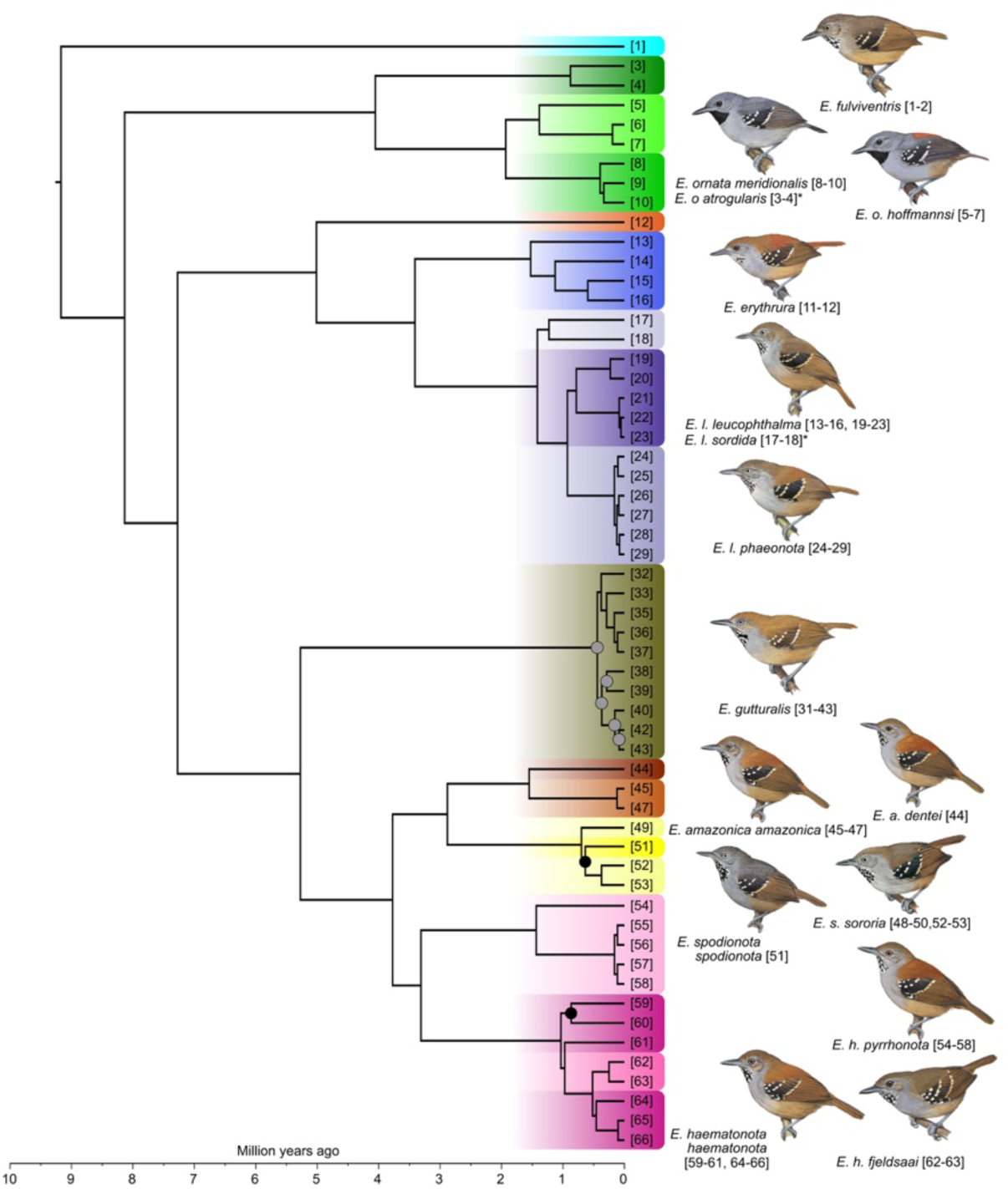
A dated phylogeny combining UCE and mitogenome sequence data. Topology estimated in Exabayes from the 75% complete phased concatenated UCE data matrix and branch lengths estimated in BEAST using the mitogenome alignments and a fixed substitution rate (see section 2.6 for details). All nodes received full support unless marked with a circle. Nodes with a gray circle with >0.75 posterior probability and nodes with a black circle with >0.90 posterior probability. A version of this tree with 95% confidence intervals on node ages is in Figure S3, and the phylogenetic tree estimated in Exabayes that contains all samples is in Figure S1A. Outgroup samples have been removed for clarity. Colors and sample numbers correspond to those in Figure 1. Illustrations (all of males) reproduced by permission of Lynx Edicions. Taxa marked with an asterisk are not illustrated and are placed below the taxon they most closely resemble in plumage.

**Figure 3.**
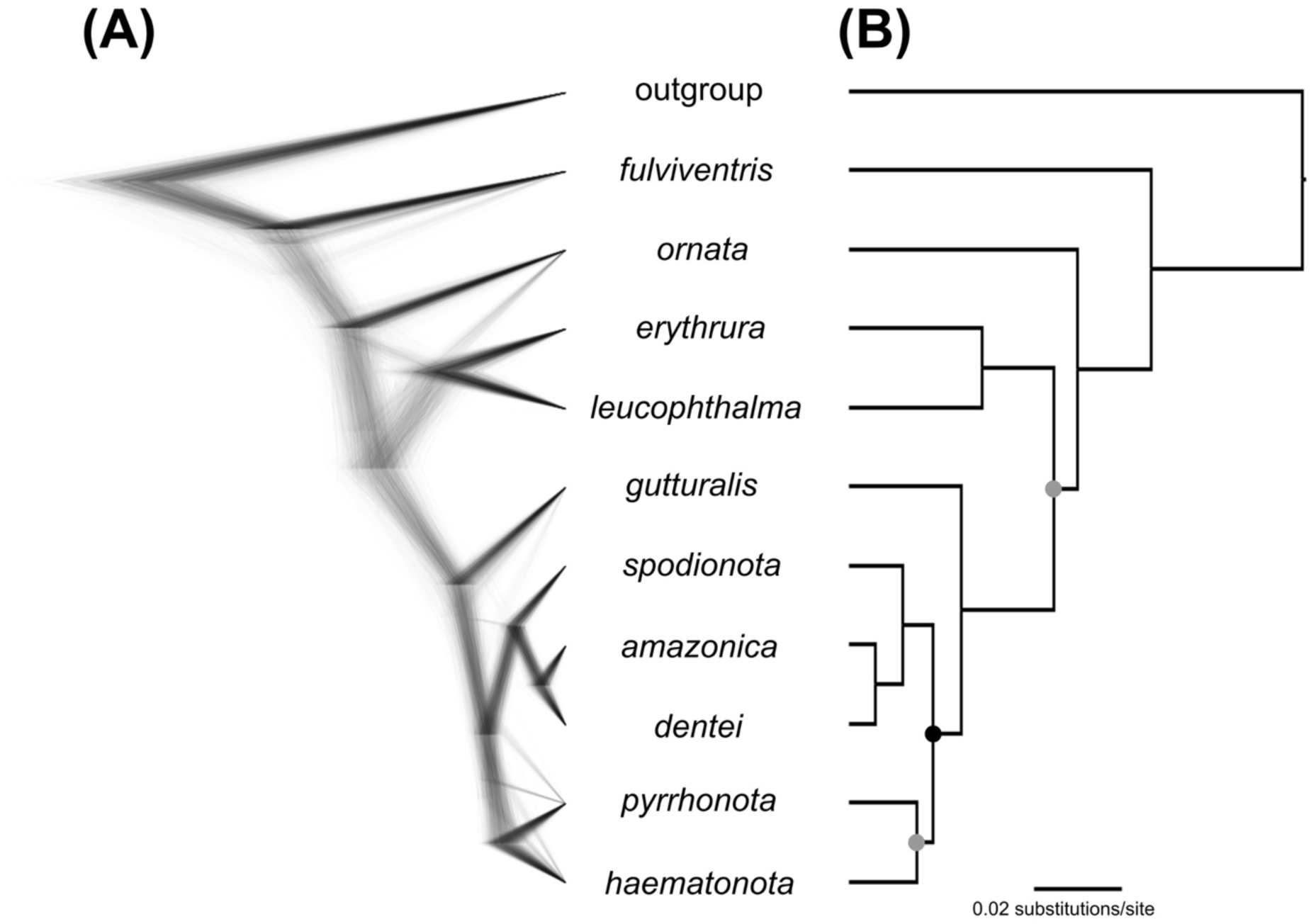
Species tree estimated in SNAPP from SNP data, using one sample per species. A) The Densitree representation of the posterior distribution of species trees and B) the Maximum Clade Credibility species tree. All nodes received full support unless marked with a circle. Nodes with a posterior probability >0.90 are marked with a black circle, and those with a posterior probability >0.75 are marked with a gray circle.

Although we lacked samples for two taxa within this species (*E. o. ornata* and *E. o. saturata*), the two parapatric gray-backed taxa occurring in Peru (*E. o. meridionalis* and *E. o. atrogularis*) are recovered as non-sister lineages, with the southern *meridionalis* sister to the rufous-backed *hoffmannsi* of eastern Brazil, and *atrogularis* sister to those two. The next split contained the sister species *E. erythrura* and *E. leucophthalma*, which together are sister to the remaining taxa (Figure 1B, Figure 2). These two species are reciprocally monophyletic, but within *E. leucophthalma* we recovered the nominate subspecies as paraphyletic. Within this nominate subspecies of *E. leucophthalma*, the western populations (samples 13–16) showed a deep divergence from the remainder of the species. The final clade contained eight parapatric taxa (*gutturalis*, *dentei*, *amazonica*, *spodionota*, *sororia*, *pyrrhonota*, *haematonota*, and *fjeldsaai*) that together range across the majority of the Amazon Basin (Figure 1C, Figure 2). The Guiana Shield taxon *E. gutturalis* was sister to the rest of the taxa in this clade, but contains minimal genetic structure in the phylogeny (Figure 2). The remaining taxa can be divided into three groups with similar divergence times between them (Figure 2). The first group contained the southeastern Amazonian *E. amazonica* (including the subspecies *dentei*) and the Andean foothill *E. spodionota* (including the subspecies *sororia*), the second is the northwestern Amazonian taxon *E. haematonota pyrrhonota*, and the third is the western Amazonian *E. haematonota haematonota* (including the subspecies *fjeldsaai*). The taxon *fjeldsaai* was embedded within *haematonota* in all analyses. Our dated phylogeny placed *pyrrhonota* sister to *haematonota*, with *amazonica* and *spodionota* sister to those. Analyses of nuclear data with a variety of phylogenetic methods (see Supplementary Material) largely supported the above results. However, some phylogenetic analyses of nuclear and mitochondrial data indicated support for two alternate topologies with regard to the placement of *E. h. pyrrhonota* (Figure 4), with varying degrees of node support (Figure S1, S2). Branch lengths in the dated phylogeny produced some results slightly discordant with those from other phylogenetic analyses, suggesting that the oldest splits within *E. ornata* and *E. leucophthalma* are as old or older than some of the species-level splits within the *haematonota s.l.* clade (Figure 2).

**Figure 4.**
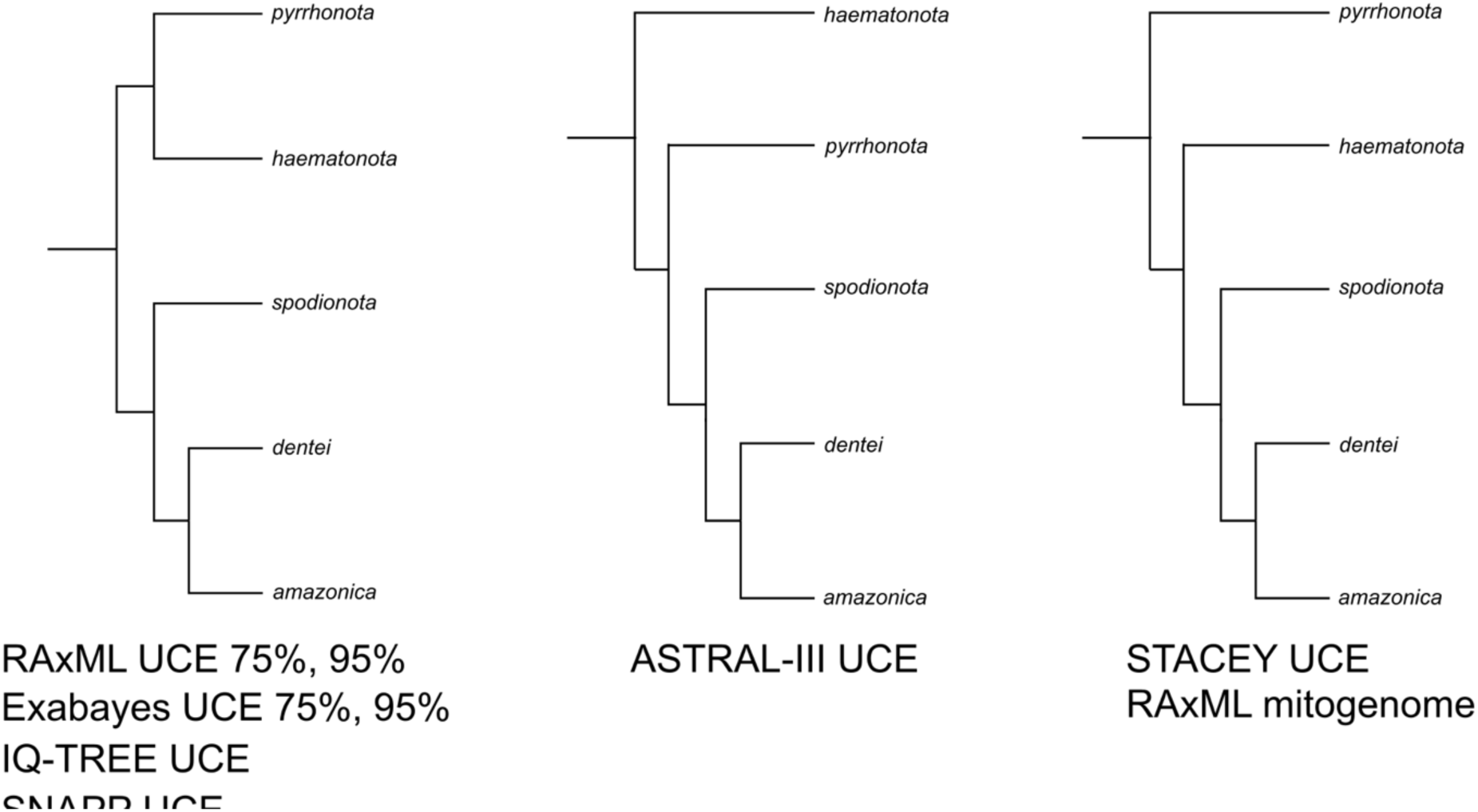
Alternate topologies recovered across phylogenetic methods for relationships within the *Epinecrophylla haematonota s.l.* clade. Results are shown using a single individual per taxon for visualization purposes. In all cases *E. haematonota fjeldsaai* was recovered as embedded within *E. h. haematonota* and is not shown. The methods recovering each topology are shown below the topology, and the full phylogenies using each method are in Figure S1.

The site-based MCC phylogeny estimated in SNAPP using one sample per species produced the same inter-specific topology as that recovered in the dated phylogeny, but with low support for two nodes (Figure 3B). The Densitree representation of the posterior of species trees showed that these nodes contained a primary topology the same as that recovered in the other nuclear analyses, but with minor alternate topologies at nodes with lower support in the MCC tree (Figure 3A). These are the same nodes that showed low support in the mitochondrial phylogeny or in which the mitochondrial phylogeny differed from the nuclear phylogeny (Figure S1, S2). The expanded SNAPP analysis using 19 samples recovered the same topology in the MCC tree, but with much lower support for multiple nodes (Figure S4B). This was reflected in considerable uncertainty in the Densitree representation of the posterior of species trees, which showed many alternate topologies both near the root of the tree and with regards to the placement of *pyrrhonota* (Figure S4A).

### 3.3. Population genetics

DAPC analyses with k-means cross-validation estimated a best fit model of K=2 for each of three clades: *E. leucophthalma*, *E. ornata*, and *haematonota s.l.*, and a model of K=1 for *E. gutturalis* (Figure 5). For *E. leucophthalma* this divided the species into the “*leucophthalma* west” clade (samples 13–16) and the remainder of the eastern taxa, including the “*leucophthalma* east” clade. For *E. ornata*, DAPC separated the central Peruvian *atrogularis* from the two eastern taxa. Lastly, for the *haematonota s.l.* group, the best fit model of K=2 separated *pyrrhonota* from the remainder of the group. The worse-fit models of K>2 (based on BIC scores) for the *haematonota s.l.* clade first separated the *E. spodionota* + *E. amazonica* clade at K=3, then *E. spodionota* from *E. amazonica* at K=4, and the western-most sample (#54) of *E. h. pyrrhonota* at K=5. PCA plots with points labeled by taxon and sample number are shown in Figure S5.

**Figure 5.**
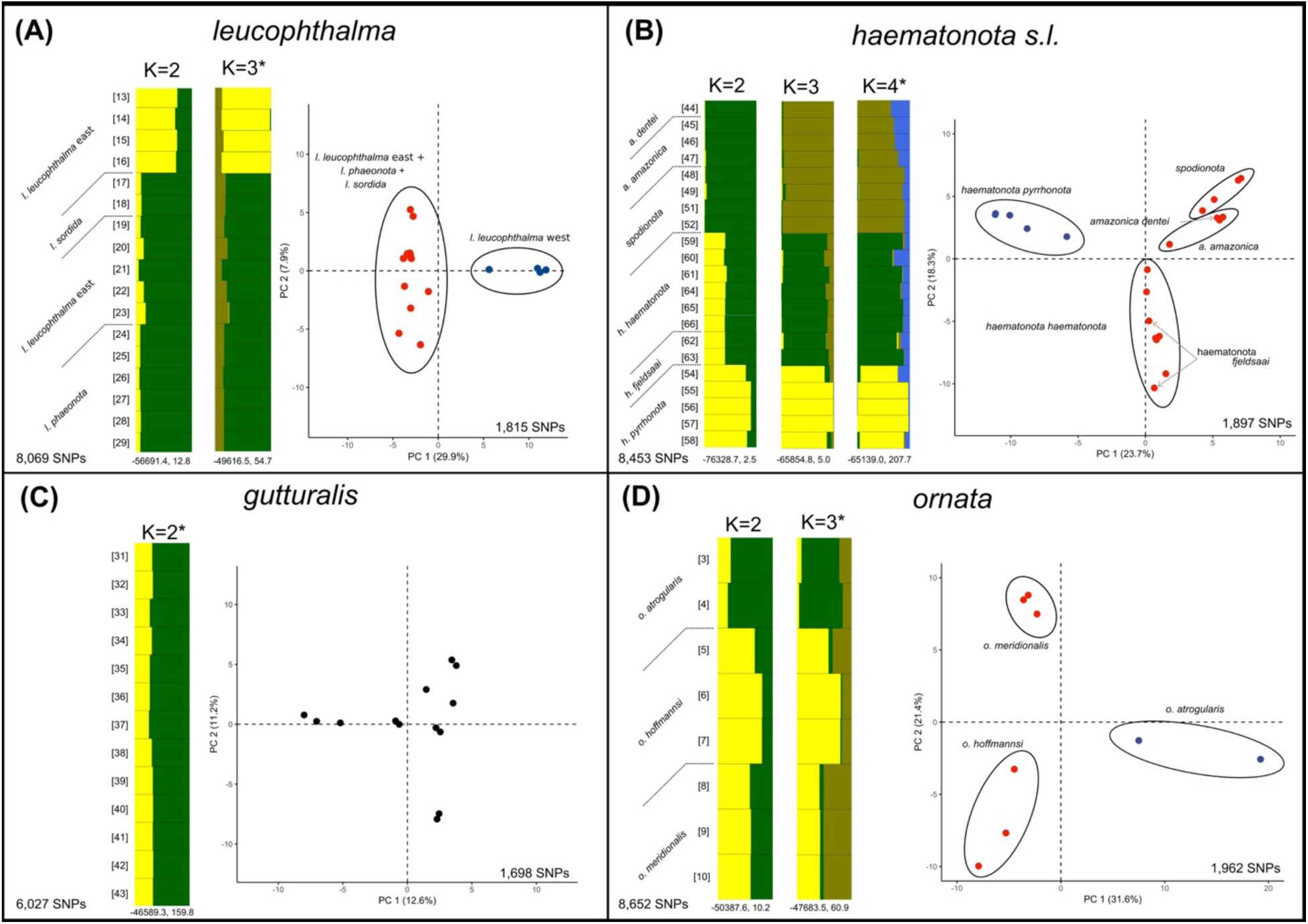
Intra-specific population genetic analyses. A) *Epinecrophylla leucophthalma,* B) the *E. haematonota s.l*. clade, containing *dentei*, *amazonica*, *spodionota*, *sororia*, *pyrrhonota*, *haematonota*, and *fjeldsaai*, C) *E. gutturalis*, and D) *E. ornata*. For each section is shown a Principal Components Analysis (PCA) with samples colored by the Discriminant Analysis of Principal Components Analysis (DAPC) group assignments on the right, and STRUCTURE plots for all likely values of K (i.e. those with low standard deviation across replicate runs) on the left. Not shown are results for K=1. The plot for the “best” value of K for each clade using the Evanno method is marked with an asterisk. Mean log likelihood and delta K values are shown below each STRUCTURE plot. Sample size in PCA plots refer to the number of unlinked SNPs recovered in that clade and used in the PCA analysis. Blue and red circles denote group assignments from DAPC while black circles and text denote taxa. Sample numbers correspond to those in Figure 1 and Table 1. PCAs with sample numbers included are shown in Figure S5.

STRUCTURE results (Figure 5) largely recapitulated those from DAPC but provided a more in-depth view of individuals with potential genetic backgrounds from multiple populations (i.e. potential introgression). Results from the Evanno method based on the ΔK value were unambiguous in all cases. However, in all cases the STRUCTURE plot for the “best” value of K from the Evanno method added a population that was evenly assigned across all individuals. Therefore, we consider the STRUCTURE plot for the “best” K minus 1 to be a more biologically realistic representation of the data and report all STRUCTURE plots >1 that have high likelihood values, following the recommendation of Meirmans (2015). Because the Evanno method is unable to calculate a ΔK value at K=1 and because all individuals of *E. gutturalis* were approximately equally assigned to both populations at K=2, we consider a K=1 to be the best-fit model for that species. For *E. leucophthalma*, K=2 separated the “*leucophthalma* west” clade from the remainder of the eastern taxa, but with all individuals containing a small percentage of genetic membership from the other clade. Within *E. ornata*, results were similar to those of DAPC, with *atrogularis* the most distinct at K=2, but with all individuals having a proportion of their ancestry assigned to both populations. The pattern in the STRUCTURE plots for the *haematonota s.l.* clade was more complex. At K=2, STRUCTURE assignments largely separated *E. h. pyrrhonota* from *E. spodionota* + *E. amazonica*, with all individuals of the *E. h. haematonota* + *E. h. fjeldsaai* group having about equal ancestry between the two groups. This pattern was also reflected also in the intermediate position of the *E. h. haematonota* + *E. h. fjeldsaai* group along the first principal component of the PCA results. At K=3 STRUCTURE separated three groups that corresponded to taxonomy, with most individuals showing only a small proportion of shared population assignments: 1) *E. h. pyrrhonota*, 2) *E. spodionota* + *E. amazonica*, and 3) *E. h. haematonota* + *E. h. fjeldsaai*. The “best” value of K=4 provided only a slight suggestion of differentiation between *E. spodionota* and *E. amazonica*, with *E. a. dentei* genetically indistinguishable from *E. a. amazonica* and *E. s. sororia* genetically indistinguishable from *E. s. spodionota*.

## 4. DISCUSSION

### 4.1. Phylogeny and population genetics

Our analyses of nuclear and mitochondrial data largely resolved the evolutionary relationships among *Epinecrophylla* taxa and recovered three broadly sympatric species complexes in the Amazon Basin. We consider the topology of the phylogeny illustrated in Figure 2 to be the best representation of relationships in the group based on the consistently high support values across multiple methods that recovered this topology (Figure 3, Figure 4, Figure S1A-D, Figure S1G). Although there was some disagreement among methods and data types regarding the placement of the taxon *pyrrhonota* (Figure 4), most of the topologies that disagreed with the sister relationship of *pyrrhonota* and *haematonota* received low support for that node, often in conjunction with a very short subtending branch (Figure S1E-F). These short branches suggest that the divergence between the three clades in the *haematonota s.l.* clade was likely very rapid, which may explain the conflicting signal across methods and the support for alternate topologies in the SNAPP posterior distributions (Figure 3A, Figure S4A). Because a strictly bifurcating tree is likely not an appropriate model for intraspecific relationships in cases of high levels of gene flow (Eckert and Carstens, 2008), our population genetics results may be a better representation of the evolutionary relationships among individuals at this fine scale. The evolutionary patterns recovered with these population genetics analyses indicate little differentiation between most subspecies and even between some taxa currently recognized as species (e.g. *E. spodionota* and *E. amazonica*). The three Amazonian species complexes that we recovered in our phylogeny are sympatric across much of the western Amazon Basin, but are represented by one species each in the eastern parts of the Basin. These species complexes are 1) *E. ornata*, 2) *E. leucophthalma* and *E. erythrura*, and 3) *E. gutturalis* and the “*haematonota s.l.*” group. Each complex contains taxa that are either allopatric or largely parapatric, with distributions typically bounded by large rivers (Figure 1). We discuss the phylogenetic results and taxonomic implications of each species separately.

### 4.2. Epinecrophylla fulviventris

*Epinecrophylla fulviventris* was recovered as sister to the remainder of the genus in all analyses and with relatively shallow divergence between our two samples in most phylogenies. We lack the geographic sampling or morphological data to make any taxonomic recommendations for this species, and thus suggest maintaining the current treatment of a monotypic *E. fulviventris* (e.g. Clements et al., 2019; Zimmer and Isler, 2003).

### 4.3. Epinecrophylla ornata

The only predominantly gray-bodied species in the genus, our results for this morphologically distinctive group are hampered by the lack of samples of the nominate taxon and the geographically adjacent taxon *saturata*. The species was described from a “Bogota” skin and is thus of uncertain provenance, although typically assumed to be from the lowlands of southern Amazonian Colombia (Peters, 1951). However, without samples from Ecuador or southern Colombia, we are unable to fully resolve the relationships within this species or recommend taxonomic changes. Despite the lack of these samples, we discovered deep splits and high population structure among all three subspecies in our phylogenetic analyses, suggesting that multiple species-level taxa occur in the group. The most genetically distinct of the three taxa in phylogenetic and population genetic analyses was *E. o. atrogularis* of central Peru, which all analyses placed as sister to *E. o. meridionalis* + *E. o. hoffmannsi*, thus contradicting the opinion of some authors (e.g. del Hoyo et al., 2019) that *hoffmannsi* represents a species distinct from the other four taxa in *E. o. ornata*. This relationship is surprising given the phenotypic similarity of *atrogularis* and *meridionalis*, which both lack the rufous back of the other three taxa in the *ornata* group and differ from each other primarily in the slightly duller underparts of female *atrogularis*. *E. o. atrogularis* and *E. o. meridionalis* potentially come into contact in the Ucayali Region of southern Peru, and further research on this contact zone is of interest given the deep genetic split between the two taxa presented here. Reports of specimens of *meridionalis* with some rufous on the back from southern Peru in Cusco and Madre de Dios have been suggested to be either variation within that taxon or evidence of introgression with one of the rufous-backed forms (Ridgely and Tudor, 1994). Based on the population genetic results (Figure 5D) presented here we suspect that the latter is a more likely explanation given that our STRUCTURE results show individuals with shared population assignments between all three subspecies that we sampled, despite the deep genetic splits among them.

### 4.4. Epinecrophylla leucophthalma and E. erythrura

Ours is the first study to suggest a sister relationship between these two species. The split between the two species is quite deep, and the two species have largely parapatric distributions (Figure 1B), but are locally sympatric in Peru (Schulenberg et al., 2007; Álvarez Alonso, 2002) without showing any morphological signs of introgression, thus confirming their status as species. Notably, we found that the nominate subspecies of *E. leucophthalma* is paraphyletic as currently recognized, with western populations of *E. leucophthalma* sister to a group containing the subspecies *sordida* and *phaeonota* and the eastern populations of the nominate subspecies. This deep genetic divergence within *E. leucophthalma* is comparable to some divergences among taxa considered to be species within the *haematonota s.l.* clade, in particular between *E. spodionota* and *E. amazonica*, but to our knowledge no morphological characters have been proposed to diagnose this western population. Excluding that western population, the remainder of the *E. leucophthalma* samples in our analysis showed extremely low divergence among them, although most of our nuclear phylogenies grouped samples into the subspecies *sordida*, *phaeonota*, and the eastern clade of *leucophthalma*. Because the geographically intermediate taxon *phaeonota* is morphologically distinctive (rufous back in *phaeonota* vs brown in all other *leucophthalma* taxa), we support the continued treatment of the three *leucophthalma* taxa as separate subspecies despite the lack of genetic divergence between them. The type locality of *E. leucophthalma* is on the right bank of the Madeira River at Salto do Jirau, Rondônia, Brazil (Pelzeln and Natterer, 1871), in the same interfluve and just 150 km to the north of our sample #20 (Figure 1B), thus suggesting that the name *leucophthalma* applies to the eastern clade of *E. l. leucophthalma* and that the Madeira River may correspond to the deep genetic break within the species. The southern Andean foothill taxon *dissita* comes into contact with our western clade of *E. l. leucophthalma* in southern Peru (Figure 1B), so the name *dissita* could potentially be expanded to include the rest of Peru and Pando, Bolivia (i.e. our “*leucophthalma* west” clade). Alternatively, a new name might be necessary for the western population of *E. l. leucophthalma*. However, genetic samples of *dissita* are needed to confirm which of these alternative treatments is appropriate. Estimates of the finite-sites mitochondrial distance and weighted nuclear F_st_ between the western clade of *E. l. leucophthalma* and all eastern populations are 6.3% and 0.29, respectively (Table S3, S4).

### 4.5. Epinecrophylla haematonota group

This clade contains eight taxa that have undergone many taxonomic rearrangements throughout their history (Cory and Hellmayr, 1924; del Hoyo et al., 2019; Dickinson and Christidis, 2014; Isler and Whitney, 2018; Peters, 1951; Remsen et al., 2019; Whitney et al., 2013; Zimmer, 1932a; Zimmer and Isler, 2003). Our phylogenetic analyses indicate that *E. gutturalis* is part of this species complex and is sister to the rest of the clade. All of our phylogenetic and STRUCTURE analyses showed no population structure within *E. gutturalis* across its range. DAPC results showed low levels of structure, but still indicated a K=1 based on BIC scores. The five samples that showed slight divergence from the main cluster in the PCA results (Figure 5C; sample numbers 32, 34, 35, 36, and 40) did not cluster based on geography. While there was some disagreement among analyses on the relationships between the rest of the taxa in this group, particularly with regard to the placement of *pyrrhonota*, most of our analyses agree with the topology in Figure 2. Following that topology, the next split in this clade presents an interesting biogeographic pattern, separating the western lowland Amazonian taxa *pyrrhonota*, *haematonota*, and *fjeldsaai*, from a clade containing the Andean foothill taxa *spodionota* and *sororia* and the southeastern Amazonian lowland *amazonica* and *dentei*. The only known sympatry between any taxa in this group occurs on the east slope of the Andes in southern Colombia where Salaman et al. (2002) reported *pyrrhonota* and *spodionota* being captured in the same mist-nets, thus necessitating at a minimum the separation of *pyrrhonota* and *E. amazonica* + *E. spodionota* as biological species. Given the similar divergence times between *pyrrhonota*, *E. amazonica* + *E. spodionota*, and *E. haematonota* + *E. fjeldsaai* we suspect that these three lineages likely represent separate biological species.

Isler et al. (1998) developed a yardstick-based system to evaluate species limits in Thamnophilidae based on vocalizations and applied this to *haematonota*, *pyrrhonota*, and *fjeldsaai*, finding that the three taxa did not differ in vocalizations and were best regarded as three subspecies of *E. haematonota* (Isler and Whitney, 2018). Our results suggest that the divergence between *pyrrhonota* and *E. amazonica* + *E. spodionota* (mtDNA 6.1%, weighted F_st_ 0.34) is comparable to that between *pyrrhonota* and *E. h. haematonota* + *E. h. fjeldsaai* (mtDNA 5.7%%, weighted F_st_ 0.25), and some analyses indicating a closer relationship between the former groups than the latter (Figure 4), albeit with weak support. This, combined with the results from our DAPC and STRUCTURE analyses which indicate that *pyrrhonota* is the most divergent taxon in this group, and in particular more so than *E. amazonica* is from *E. spodionota*, that *E. pyrrhonota* is best regarded as a distinct species. A more thorough evaluation of the utility of this yardstick system between all *Epinecrophylla* taxa is desirable, although the results presented here indicate that it may not be a reliable indicator of species status in all cases.

Our results support the continued treatment of *fjeldsaai* as a subspecies of *E. haematonota* due its morphological distinctiveness, but were more ambiguous with regard to the taxonomic status of *dentei*. None of our analyses were able to differentiate *fjeldsaai* from *haematonota*, and all phylogenetic analyses indicated that *fjeldsaai* was embedded within *haematonota*. This treatment is further supported by evidence of hybridization between the two taxa in northwestern Peru (LSUMNS specimens). In fact, one of our samples of *fjeldsaai* (sample 62) has some rufous coloration on the lower back suggestive of hybridization. *E.* a. *dentei* was placed as sister to *E. a. amazonica* in all phylogenetic analyses, but with relatively shallow divergence (mtDNA 2.8%, weighted F_st_ 0.19). Our population genetic analyses including all samples in the *haematonota s.l.* clade were unable to distinguish *amazonica* and *dentei* (Figure 5), and in fact were largely unable to differentiate *E. amazonica* and *E. spodionota*. A DAPC analysis using just the three samples of *amazonica* and the one of *dentei* did suggest a K=2 was the best model based on BIC scores and separated the *dentei* sample from the rest of *amazonica* (Figure S6).

In summary, we recommend the following 4-species treatment for the taxa in the *haematonota* group: *E. haematonota* (with *fjeldsaai* as a subspecies), *E. pyrrhonota*, *E. amazonica* (with *dentei* as a subspecies), and *E. spodionota* (with *sororia* as a subspecies). Until genetic samples of *E. leucophthalma dissita*, *E. o. ornata*, and *E. o. saturata* are available for study, we refrain from making taxonomic recommendations in those groups, but we suspect that both *E. leucophthalma* and *E. ornata* contain multiple species-level taxa. Therefore, we recommend the following species-level linear taxonomy for *Epinecrophylla*: *fulviventris*, *ornata*, *erythrura*, *leucophthalma*, *gutturalis*, *haematonota*, *pyrrhonota*, *amazonica*, *spodionota*.

### 4.6. Biogeographic patterns

Having three broadly sympatric species or species complexes distributed across the Amazon Basin provides replicated evolutionary histories across a shared landscape. Of interest is the response of each of these species or species complexes to well-known biogeographic barriers in the Amazon Basin, such as large rivers (Capparella, 1991; Wallace, 1854). The major river systems of the Amazon Basin, such as the Solimões, Negro, and Madeira all appear to have an effect on the genetic structure and range limits of *Epinecrophylla* taxa, delimiting many species and subspecies that show significant genetic breaks at those locations in our analyses.

Smaller river systems, however, appear to have idiosyncratic effects on genetic structure, with some delimiting genetic groups in one species, but having little to no effect in others. For example, the Purús River is a major barrier for *E. ornata*, but has little effect on the genetic structure of other groups, while the Tapajos River is a barrier for *E. leucophthalma* but not *E. ornata*. Additionally, the distribution and genetic boundaries of phenotypically distinctive taxa such as *fjeldsaai* and *phaeonota* do not appear to always follow biogeographic barriers that affect other bird species. The brown-backed *fjeldsaai*, which we find to be phylogenetically embedded within the rufous-backed *haematonota*, hybridizes with *haematonota* within the Napo interfluve without any clear biogeographic barrier separating the two taxa. Likewise, the rufous-backed *phaeonota* is part of the otherwise brown-backed *leucophthalma* group, and appears to replace the eastern populations of nominate *leucophthalma* somewhere between the Juruena and Roosevelt Rivers.

Although all three *Epinecrophylla* species complexes overlap in much of the western Amazon Basin, there is evidence that competition may play a role in their ability to coexist over broad spatial scales. For example, J. Tobias (personal communication) found *E. leucophthalma* and *E. amazonica* regularly at the same site in Madre de Dios, Peru, but the two species rarely occurred in the same mixed species flock. Likewise, in Napo, Ecuador, Whitney (1994) found *E. ornata* and *E. erythrura* in the same mixed species flock on just one occasion, although three species of *Epinecrophylla* occurred at the site.

That two species complexes – *E. ornata* and *E. leucophthalma* + *E. erythrura* – are absent from the Guiana Shield and the northern half of the Inambari interfluve (Figure 1A, 1B) is perplexing. This pattern may be due to the vagaries of extinction, interspecific competition, or habitat suitability, or some combination of those factors, all of which require further study. For example, suboptimal habitat may increase competition between such closely related and ecologically similar species, leading to local extinctions. Alternatively, more drastic climate fluctuations in the eastern half of the basin may be driving this pattern. There is evidence that the eastern Amazon Basin became much drier during the glacial periods of the Pleistocene, whereas climate of the western Amazon Basin has maintained more stable over the same time period, which is thought to have resulted in relatively higher losses of biodiversity in the eastern half of the basin (Cheng et al., 2013). A north-south gradient in vegetation composition exists in the Inambari interfluve (Tuomisto et al., 2019) and the Guiana Shield is relatively drier than much of the western Amazon (Fick and Hijmans, 2017), so these species may be unable to persist in these areas. Likewise, the *haematonota* group is absent from the Brazilian Shield, while the other two species complexes are present there.

### 4.7. Areas of potential future research

The results of our phylogenetic and population genetic analyses suggest that multiple geographic regions could produce valuable insights with both greater sampling effort and natural history observations (e.g. playback experiments, surveys of contact zones, analysis of vocal and morphological traits). The first is in southern Peru in the foothills of southeastern Madre de Dios region, where three taxa potentially come into close geographic proximity, namely *spodionota* and *amazonica*, which we recover as sister taxa in our phylogenies, and *haematonota*. A second region of interest is slightly to the north in southern Ucayali region, where two subspecies of *ornata*; *atrogularis* and *meridionalis*, replace each other, perhaps across the Purús River. These two taxa are recovered as non-sister taxa in our phylogenies and could perhaps come into contact across the headwaters of that river. Genetic samples of the two northern taxa in the *ornata* group, including the type taxon, are critical to resolving relationships within that clade. A third region is the headwaters of the Rio Napo in northern Ecuador, where two taxa currently regarded as subspecies of *haematonota*; *pyrrhonota* and *fjeldsaai*, could potentially come into contact. Analysis of a contact zone in this region would be critical to resolving species limits in the *haematonota* group.

Despite being the most well-sampled phylogenetic study of *Epinecrophylla* to date, our study lacked genetic samples from some key areas that could affect the results presented here. The lack of samples for two subspecies of *E. ornata*, including the nominate, hinders our ability to make any taxonomic recommendations for that species. We also lack samples of *E. leucophthalma dissita* of the Yungas. This taxon comes into contact with the western clade of nominate *leucophthalma*, and it is possible that the name *dissita* could apply to the entirety of this clade of western *leucophthalma*, a population that based on our results may represent a species distinct from the eastern three taxa in the *leucophthalma* group. It is worth noting that additional taxa have been named within *E. leucophthalma*, but the validity of those taxa has been questioned as they are generally considered not morphologically diagnosable (del Hoyo et al., 2019). Two other sampling gaps bear mention; the first is the population of *E. h. haematonota* from the north bank of the Amazon west of the Napo, which is the population that presumably intergrades with *E. h. fjeldsaai*, and the second is a lack of samples for any taxon from the vast region of the Brazilian Amazon west of the Madeira River and south of the Amazon River, which could contain genetically distinct populations and contains the type locality of *E. amazonica* (Peters, 1951).

## 5. Conclusions

As has been shown in other Neotropical avian systems (Brumfield, 2005; Musher and Cracraft, 2018), our study highlights the importance of sampling populations below the species level, especially in tropical regions, where the taxonomy of many groups is unresolved and there may be considerable undiscovered morphological and genetic diversity. Our understanding of phylogenetic relationships has grown dramatically in recent decades as technological advances have allowed us to obtain and analyze sequence data for ever more genetic markers and individuals, including at the population level in non-model organisms (Harris et al., 2018; Tan et al., 2019; Zarza et al., 2016; Zucker et al., 2016).

## Acknowledgements

OJ was supported by the National Science Foundation Graduate Research Fellowship under grant no. DGE-1247192. JTH was supported by the National Science Foundation Research Experiences for Undergraduates under grant no. DEB-1146265. We thank the curators and staff at the following institutions for providing tissue samples: Gary Graves and Christopher Milensky at the Smithsonian National Museum of Natural History, Richard Prum and Kristof Zyskowski at the Yale Peabody Museum, Christopher Witt at the Museum of Southwestern Biology, Joel Cracraft and Paul Sweet at the American Museum of Natural History, John Bates and Ben Marks at the Field Museum of Natural History, and Rob Moyle and Mark Robbins at the University of Kansas Biodiversity Institute & Natural History Museum. Donna Dittmann assisted with tissue sample processing at the Louisiana State University Museum of Natural Science. Van Remsen, Daniel F. Lane, Nelson Buainain Neto, and members of the Brumfield Lab provided invaluable feedback on versions of this manuscript. Marco A. Rego provided assistance with the creation of Figure 1.

## SUPPLEMENTAL CAPTIONS

Supplemental Figure 1. Seven estimates of the phylogeny of *Epinecrophylla*, based on UCE alignments, showing congruence of internal topologies across methods. A) Exabayes tree estimated from the 75% complete concatenated data matrix (note that this tree is also shown in Figure 2), with node support values of posterior probability. B) Exabayes tree estimated from the 95% complete concatenated data matrix, with node support values of posterior probability. C) RAxML tree estimated from the 75% complete concatenated data matrix, with node support values of bootstrap likelihood. D) RAxML tree estimated from the 95% complete concatenated data matrix, with node support values of bootstrap likelihood. E) STACEY tree estimated from the fully-partitioned alignment using the 150 loci containing the greatest number of parsimony-informative sites, with node support values of posterior probability. F) ASTRAL-III tree estimated from RAxML gene trees using the 75% complete data matrix, with node quartet support values of local posterior probability. G) IQ-TREE estimated from the 75% complete data matrix. Node support values shown in ultrafast bootstrap likelihood, gene tree concordance factors, and site concordance factors, respectively.

Supplemental Figure 2. An estimate of the phylogeny of *Epinecrophylla*, based on an alignment of draft mitochondrial genomes using a Maximum Likelihood methodology implemented in RAxML, with node support values of bootstrap likelihood.

Supplemental Figure 3. The dated phylogeny shown in Figure 1, with nodes showing the 95% highest posterior density of the divergence estimates based on 11,488 base pairs of the draft mitochondrial genomes and a fixed mitochondrial substitution rate.

Supplemental Figure 4. Species tree estimated in SNAPP from SNP data, using 1-2 samples per clade. A) The Densitree representation of the posterior distribution of species trees and B) the Maximum Clade Credibility species tree. All nodes received full support unless marked with a circle. Nodes with a posterior probability between 0.90 and 0.75 are marked with a gray circle and those <0.75 are marked with a white circle. No nodes received a posterior probability between 1 and 0.90. See Figure 3 for the SNAPP tree using one sample per species.

Supplemental Figure 5. PCA plots shown in Figure 5 with samples labeled by taxon and sample number.

Supplemental Figure 6. PCA plot using four samples of *Epinecrophylla amazonica*. DAPC results indicated a best fit model of K=2, separating the one sample of *dentei* from the three of *amazonica*.

Supplemental Table 1. Samples removed from analyses due to misidentification or potential contamination.

Supplemental Table 2. Samples for which we were unable to recover mitochondrial genomes either due to a failure with MITOBIM or very high amounts of missing data in the recovered mitochondrial genome.

Supplemental Table 3. Matrix of mitochondrial genetic distances between taxa in this study. Values above the diagonal represent the uncorrected *p*-distance and those below the diagonal represent the genetic distance accounting for multiple substitutions under the GTR + γ + I finite-sites substitution model. Values in the diagonal represent the average intra-taxon distance for the finite-sites distance (left) and the uncorrected *p*-distance (right). Both distance methods are based on the concatenated alignment of the 13 mitochondrial protein coding genes. All values are shown as a percentage.

Supplemental Table 4. Matrix of nuclear genetic distances between taxa in this study. Values below the diagonal are the weighted value of F_st_ between taxa, averaged across UCE loci. Values in the diagonal represent the average intra-taxon distance.

Supplemental Table 5. Individuals used as references for SNP calling.

